# DNA methylation-based immune cell deconvolution in solid tumors

**DOI:** 10.1101/619965

**Authors:** Cong Liang, Xiaoqing Yu, Bo Li, Y. Ann Chen, Jose R. Conejo-Garcia, Xuefeng Wang

## Abstract

Understanding of the tumor microenvironment (TME) structure is likely to have a profound and immediate impact on therapeutic interventions as well as the development of signatures for diagnostic and prognostic evaluations. DNA methylation arrays represent one of the most reproducible molecular assays across replicates and studies, but its value of profiling tumor-infiltrating immune lymphocytes (TILs) hasn’t been intensively investigated. Here we report a model-based evaluation of tumor TIL levels using DNA methylation profiles. By employing a hybrid method of stability selection and elastic net, we show that methylation array data in ten TCGA cancer types provide a strikingly accurate prediction of immune cell abundance, in particular the levels of T cells, B cells and cytotoxic cells in skin cutaneous melanoma (SKCM). The immune-informative CpG sites showed significant prognostic values, representing important candidates for further functional validation. Further, we present regression models each using only ten CpG sites to estimate the levels of infiltrated immune cell types in melanoma. To validate these models, we performed matched methylation EPIC array and RNA-seq on 30 new melanoma samples. We observed high concordance on methylation and gene expression predicted tumor immune infiltration levels in our new dataset. Our study demonstrated that DNA methylation data is a valuable resource in reliably evaluating tumor immune responses. The selected methylation panels provide candidate targets for future clinical researches. Our prediction models are easy to implement and will provide reference for future clinical practices.

## Introduction

Understanding of the crosstalk between tumor cells and the host immune microenvironment is crucial to the prediction and monitoring of therapy response, and to the discovery of new targets for drug development [1]. In addition to higher mutation and neo-antigen load [2–4], the presence of tumor infiltrating lymphocytes (TIL) is believed to be associated with a favorable prognosis and better response to adjuvant treatment. For example, gene expression signatures specific to CD8+ cytotoxic lymphocytes and dendritic cells have been found to be associated with a better overall clinical outcome in cancer [5]. Insights on the roles of these immune cell types in cancer progression and immune evasion, as well as the association between other immune cells and drug responses, offer new opportunities for more effective interventions[1, 6, 7].

However, there are still considerable technological and analytical barriers to accurately assessing tumor immunity in situ. The major disadvantages of traditional H&E and immunohistochemical (IHC) staining methods are that they are only semi-quantitative and that they suffer from bias and variability from sample slicing. Flow cytometry analysis offers more accurate immune cell measures, but it is labor intensive and requires fresh tissues and cell type specific markers. Over the past decade, efforts has been made to deconvolve the tumor microenvironment (TME) from microarray or RNA-seq profiled gene expression data. CIBERSORT applies a support vector regression of tumor gene expression profiles on a matrix of reference gene expression signatures [8]. TIMER is a resource that employs a constrained linear regression model on expression levels of informative genes [5]. Both of these two methods require reference gene expression profiles from purified immune cells to identify informative signatures and perform the estimation. On the other hand, single sample gene set enrichment analysis (ssGSEA) calculates the expression enrichment score for a predefined marker gene list within a sample [ref]. Gene expression profiles in reference cells are not required in using ssGSEA, which avoids the potential bias introduced by references. Molecular researches have provide plenty of resources for marker genes in various immune cells. Some are summarized in Bindea et. al. [9], as well as in the latest nCounter PanCancer Immune profiling panel from NanoString [10]. Studies using the above methods have observed associations between tumor immune infiltration and cancer prognosis in multiple cancer types, which offers great promise in their clinical applications [11, 12]. However, significant variations have been observed between estimations using different methods or references. There is still a pressing need to improve the accuracy in quantifying the cell components of TME to facilitate both retrospective and prospective clinical studies.

DNA methylation has an essential role in the epigenetic control of gene expression and disease development. The haploid human genome has approximately 29 million cytosine-Guanine (CpG) sites with different methylation status [13], which is collectively referred to as the DNA methylome. An increasing number of cancer methylome profiles from tumor and other tissues have been accumulated in the public domain. To date, TCGA has processed >10,000 samples with two types of array platforms, Infinium HumanMethylation27 (27k array, released in 2009) and HumanMethylation450 (450k array, released in 2011) [14]. They measure around 27,000 and 485,000 individual CpG sites respectively. More cancer samples have been now profiled with HumanMethylationEPIC array (contains over 860k sites) and other platform such as bisulfite sequencing. The methylome profiles have been found to provide stable cell differentiation signatures [ref], and studies have found that it can accurately estimate cell components in blood samples [15]. Recently, the reference-based method CIBERSORT has been employed to estimate the cellular composition of 9 cell types from DNA methylation data [16]. In addition, methylation assay have much less stringent tissue sample requirement: it requires small amount of DNA and does not require fresh tissues. As such, there is a strong impetus for a comprehensive analysis of tumor methylomes for immune cell type and response deconvolution. The predictive biomarker panels based on DNA methylation have great translational potential due to its high detection sensitivity and stable signatures.

In this study, we proposed a model-based method to evaluate the level of tumor infiltrated lymphocytes. We took advantage of the tumor immune cell infiltration results from gene expression profiles using ssGSEA and TIMER. By applying a hybrid method of stability selection and random lasso to 10 cancer types in TCGA, we selected CpG sites (methylation panel CpGs) that are important in predicting tumor immune cell scores for each of 24 immune cell types. We found that genes in close proximity of methylation panel CpGs are enriched in immune response related functions. We observed associations between methylation of panel CpG sites and cancer prognosis. Finally, we focused on three cell types: T cells, B cells, cytotoxic cells in skin cancer, which achieved the best prediction accuracy in the TCGA dataset. We arrived at a simple linear regression model with ten variables (LS10) for each of the three cell types, and validated our models with 30 newly sequenced skin cancer samples with matched DNA methylation array and RNA-seq profiles. Our study provides a facile model for evaluating levels of immune infiltration in skin cancer using DNA methylation data. The cell type and cancer type specific methylation panels will also serve as an important resource for future clinical studies.

## MATERIAL AND METHODS

### TCGA methylation and gene expression data

We analyzed 10 major cancer types each with at least 300 samples that have matched methylation profiles and gene expression profiles available in TCGA: bladder urothelial carcinoma (BLCA), breast invasive carcinoma (BRCA), head and neck squamous cell carcinoma (HNSC), kidney renal clear cell carcinoma (KIRC), brain lower grade glioma (LGG), lung adenocarcinoma (LUAD), lung squamous cell carcinoma (LUSC), prostate adenocarcinoma (PRAD), and skin cutaneous melanoma (SKCM) and thyroid carcinoma (THCA). Molecular profiles for all tumor samples were downloaded from the Broad’s GDAC Firehose portal (http://gdac.broadinstitute.org/). Normalized beta values from the Illumina Infinium HumanMethylation450 platform (450K) containing 485,577 probes were downloaded. CpG sites with more than 5% missing values were removed from downstream analysis and the rest of missing values were filled by the median of available data. CpG sites with bottom 10% variances were also removed to minimize the influence of observational noise. We performed an arcsine transformation of the beta values: arcsin(2*Beta-1) before applying further statistical analysis [ref]. Normalized gene expression values from RNA-seq were also downloaded for matched samples. We used two published methods to estimate the level of infiltrated immune cells in this dataset. 1) Tumor Immune Estimation Resource (TIMER, http://cistrome.org/TIMER) estimates the relative abundance of the six major tumor-infiltrating immune cell types, CD8+ T cell, CD4+ T cells, B cells, neutrophils, macrophages, and dendritic cells [5]. 2) Single cell gene set enrichment analysis (ssGSEA) estimates the level of infiltration for 24 immune cells, 2 lymphocyte infiltration summary statistics, as well as the activity of tumor antigen presenting machinery. 24 immune cell type specific signature genes were adopted from Bindea et. al. [9].

Two lymphocyte infiltration summary statistics are defined as in Senbabaoglu et. al. [12]: (1) overall immune infiltration score (IIS) is aggregated from both adaptive and innate immune scores; (2) T cell infiltration score (TIS) is aggregated from nine T cell scores. ssGSEA scores were calculated using R bioconductor package GSVA [17].

### Hybrid method to get methylation panel

We utilized an iterative method as shown in Figure 2A to identify marker CpG sites in tumor immune infiltration. In each iteration, ⅔ of all samples were randomly selected as training set, while the rest of samples were kept as test set. Next, 10% features were randomly selected as candidate features for the regression analysis. Then elastic net regression were performed using R package *glmnet* [18]. This process was iterated 500 times in consideration of execution time and stability of the results. At the end of all iterations, importance score of each CpG were calculated, and the prediction accuracy was summarized. The importance score is defined as the ratio of the counts that a feature is kept by elastic net divided by the counts that this feature is randomly chosen in the feature pre-selection step. Features with importance score greater than 0.9 (methylation panel CpG sites) were documented in the methylation panel in Supplementary table 2. R source code for feature screening and selection is available at https://github.com/xfwang/immu.

**Figure 1.**
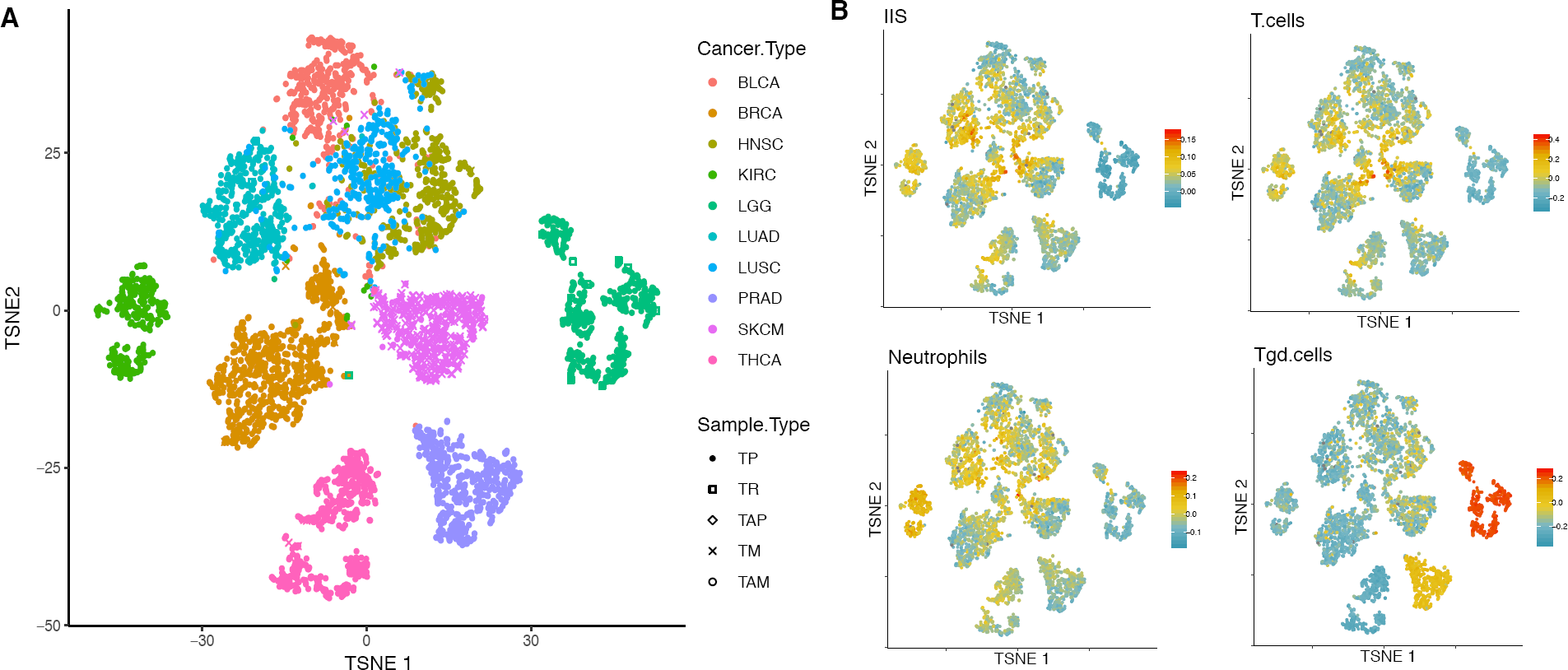
Global methylation profiles across cancer types. **(A)** Visualization of methylation profiles of all TCGA samples in this analysis using t-SNE. TCGA sample type codes: TP - primary solid tumor; TR - Recurrent solid tumor; TAP – additional, new primary; TM – metastatic; TAM - additional metastatic. **(B)** Visualization of selected immune scores of all samples. Immune scores were calculated using ssGSEA method.

**Figure 2.**
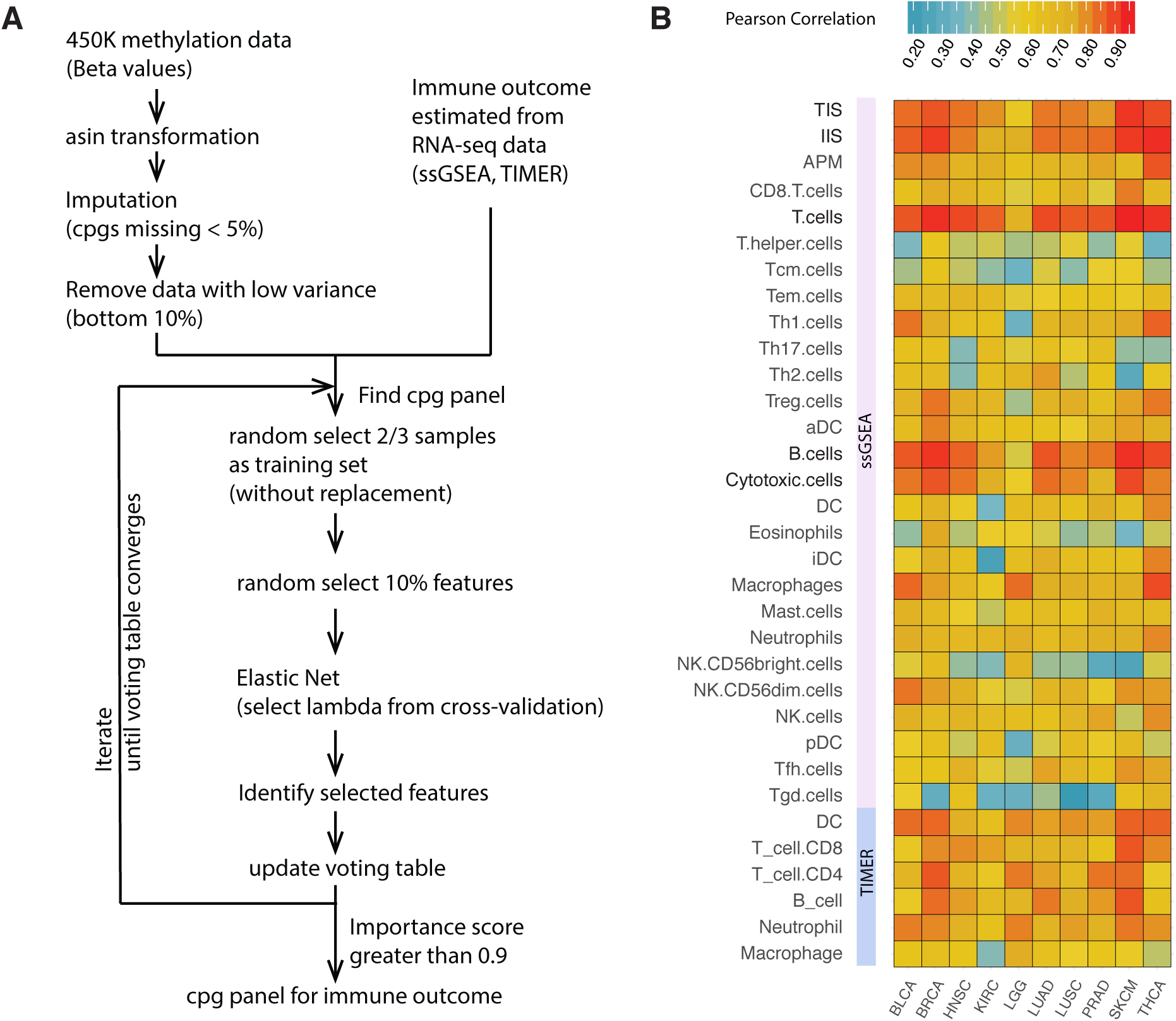
A hybrid method to evaluate levels of tumor infiltrated immune cells. **(A)** The workflow to derive methylation panels. **(B)** DNA methylation profiles is powerful in estimating tumor infiltrated immune cell scores. Prediction accuracy varies by cancer and immune cell type. In general, skin cancer melanoma (SKCM) achieves the best prediction accuracy among cancer types. T cells, B cells as well as cytotoxic cells achieves the best prediction accuracy among cell types.

### Gene ontology analysis

Gene ontology analysis was performed using Gorilla, using all human protein coding genes as background [19].

### Survival analysis

Clinical data for TCGA dataset were downloaded from https://www.cbioportal.org/. We evaluate the association of each individual CpG biomarker in the methylation panels with patient overall survival (OS) and progression-free survival (PFS) based on Kaplan-Meier method and log-rank test. The log-rank test p values were calculated using R package *survival* [20] The survival plots were generated using R package *survminer* and *ggplot2* [21].

### SKCM EPIC array and RNA-seq data analysis

R package *minfi* was used to preprocess the 850K methylation data from EPIC array [22]. CpGs with less than 5% missing value were imputed by the median value of the probe. Quantile normalization method was applied to get beta values. RNA-seq data were aligned to hg19 using TopHat2. Reads were counted using htseq-count. Normalized gene expression TPMs were calculated using RSEM. ssGSEA scores were calculated for 30 new SKCM samples the same as TCGA samples.

## RESULTS

### Global methylation profiles across tumor types

We analyzed 10 major cancer types that have more than 300 samples with both 450K methylation profiles and RNA-seq gene expression profiles available in TCGA. The data include 407 bladder urothelial carcinoma (BLCA), 780 breast invasive carcinoma (BRCA), 521 head and neck squamous cell carcinoma (HNSC), 316 kidney renal clear cell carcinoma (KIRC), 530 brain lower grade glioma (LGG), 449 lung adenocarcinoma (LUAD), 370 lung squamous cell carcinoma (LUSC), 498 prostate adenocarcinoma (PRAD), 471 skin cutaneous melanoma (SKCM) and 509 thyroid carcinoma (THCA) samples. We performed an unsupervised clustering analysis of their methylation profiles using the t-distributed stochastic neighbor embedding (t-SNE) method. As shown in Figure 1A, patient samples predominantly cluster by cancer types. Clusters from squamous histology cancers LUSC and HNSC partly overlap each other. Some cancer subtypes such as basal-like breast cancers also make up distinguished clusters (Supplementary Figure 1). We calculated tumor infiltrating lymphocyte (TIL) scores from RNA-seq data using two methods, TIMER and ssGSEA, (see methods) and mapped the results to the t-SNE plot. We found the level of immune cell infiltration varies among cancer and immune cell types (Supplementary Figure 2). For example, compared to other cancer types, an immune “cold” cancer type, brain lower grade glioma, has lower abundance for most immune cells but the highest gamma delta T cells (Tgd cells) abundance (Figure 1B-E). These results suggest that the methylation profiles for cancer samples serve as an important molecular signature for their states and may provide important information for cancer treatment and prognosis.

### Estimate tumor immune landscape using DNA methylation profiles

We applied a supervised learning method to estimate TIL scores and to identify CpG biomarkers that are important in estimating tumor immune landscape (Figure 2A, see methods). The TIL scores estimated from RNA-seq using ssGSEA were used as the training response of the immune infiltration level. Compared to gene expression, DNA methylation data are ultra-high dimensional, thus standard feature selection methods such as lasso and elastic net cannot be directly applied due to high computational complexity and unstable estimation. To solve this issue, we employed a hybrid method that combine the advantages of stability selection and random lasso. Stability selection method generates multiple bootstrap samples from the original data, and increases the stability of the result by summarizing the results from multiple bootstraps [23]. Random lasso keeps a subset of features for learning in each iteration, thus alleviate the collinearity issue in the data [24]. Our method is based on an iterative process with the following steps: Samples were first randomly split into training and test set in 2:1 ratio. 10% CpG sites were randomly selected as candidate features before the elastic net model fitting. Using elastic net regression, features were further selected because a fraction of coefficients were suppressed to zero due to L1 regularization. The regularization parameter in elastic net was chosen based on a cross-validation process. We next evaluated the prediction accuracy using test set Pearson and Spearman correlations as well as mean squared error. These steps were iterated for 500 times, considering both the computational complexity and the result stability. Finally, we defined an importance score of each CpG, as its ratio of non-zero coefficients in training models where the CpG is a candidate feature. The CpGs were prioritized by their importance scores. Importance score = 1 indicates the feature is an important signature in predicting TIL levels. Whereas importance score = 0 indicates that the feature is not important in predicting TIL levels.

We observed a high concordance between methylation and RNA-seq predicted TIL scores for 24 cell types and 3 summarizing scores using ssGSEA (as defined in [12], see method, Figure 2B and Supplementary Table 1). We found that methylation data can accurately predict overall immune infiltration level. The Pearson correlation between methylation and ssGSEA estimated immune infiltration score (IIS) ranges from 0.75 to 0.93. The Pearson correlation for T cell infiltration score (TIS) varies from 0.59 to 0.92, with most cancer types exceed 0.8 expect for LGG (0.59). In predicting specific immune cell types, we found that methylation data achieved the highest prediction power for T cells (Pearson correlation ranges from 0.86 to 0.95) and B cells (Pearson correlation ranges from 0.78 to 0.93). Cytotoxic cells, estimated from ssGSEA, also achieved a high prediction concordance between methylation data and gene expression data (Pearson correlation ranges from 0.57 to 0.90). As expected, cell types with lower abundance in the tumor microenvironment, such as T gamma delta (Tgd) cells and eosinophils, are more difficult to predict. In terms of cancer types, SKCM and BRCA have higher prediction accuracy compared to other cancer types. In contrast, LGG obtains low prediction accuracy in general. In addition, we applied our methods to TIL scores for six immune cell types estimated in TIMER (Figure 2B). We observed high Pearson correlation between methylation and gene expression predicted TIL scores for five out of six major immune cell types: CD4 T cells (varies from 0.58 to 0.87), CD8 T cells (from 0.59 to 0.87), B cells (from 0.59 to 0.88), dendritic cells (from 0.65 to 0.86) and neutrophils (from 0.63 to 0.83). The prediction accuracy for macrophage is lower compared to other cell types (Pearson correlation ranges between 0.39 and 0.76). With respect to cancer types, SKCM and BRCA again achieved the best prediction accuracy, with Pearson correlation ranges from 0.58 to 0.88.

### Important predictive and prognostic methylation biomarkers

We further investigated specific CpG biomarkers in estimating T cells and B cells, focusing on five cancer types (BRCA, HNSC, LUAD, PRAD and SKCM) with a high prediction accuracy. Top ranking CpGs are methylation markers that obtain a high predictive power of the tumor immune microenvironment. We first investigated top CpGs that are shared by different cancer types. The importance scores of three CpG sites, cg04776231, cg14094409, and cg04776231 are greater than 0.9 for T cell in all five cancer types (Figure 3A). These CpGs are in the gene body of PTPN12, DIABLO, and CCDC57. PTPN12 is a member of protein tyrosine phosphatase (PTP) family gene which has been identified as an important prognosis marker in multiple cancer types. Previous study has observed abnormally low PTPN12 expression in triple negative breast cancer patients, while restoring PTPN12 expression significantly impacted the tumorigenic and metastatic potential of PTPN12 deficient cells [25]. Decreased expression of PTPN12 is also correlated with poor prognosis in hepatocellular carcinoma [26] and non-small cell lung cancer [27]. *Diablo* (also called *smac*) is a protein that interacts and antagonizes inhibitors of apoptosis proteins (IAPs) [28], and it has been identified as a prognosis marker in multiple cancers like colon cancer [29], small cell lung cancer [30]. In the case of estimating B cell abundance, three CpG sites, cg26568226, cg01445100 and cg15286847, have importance scores greater than 0.9 in our analysis (Figure 3B). They are located in the gene body of CYFIP1, BANP, and KLHL36, respectively. *Cyfip1* is a component of WAVE regulatory complex that promotes actin assembly. It was found to be commonly deleted in various human epithelial cancers, and it may serve as an invasion suppressor gene [31]. BANP encodes a protein that binds to matrix attachment regions, but its gene expression level doesn’t significantly vary among tissue types according to GTEx results from UCSC genome browser. BANP gene can also generate a circular RNA from its exon 5-11 (circ-BANP). Circ-BANP has been found to be overexpressed in colorectal cancer and lung cancer samples, and was suggested as a prognosis and therapeutic marker for colorectal and lung cancer [32, 33]. KLHL36 is a gene that was much less investigated in cancer cells compared to the other two, and its molecular functions still remain investigated.

**Figure 3.**
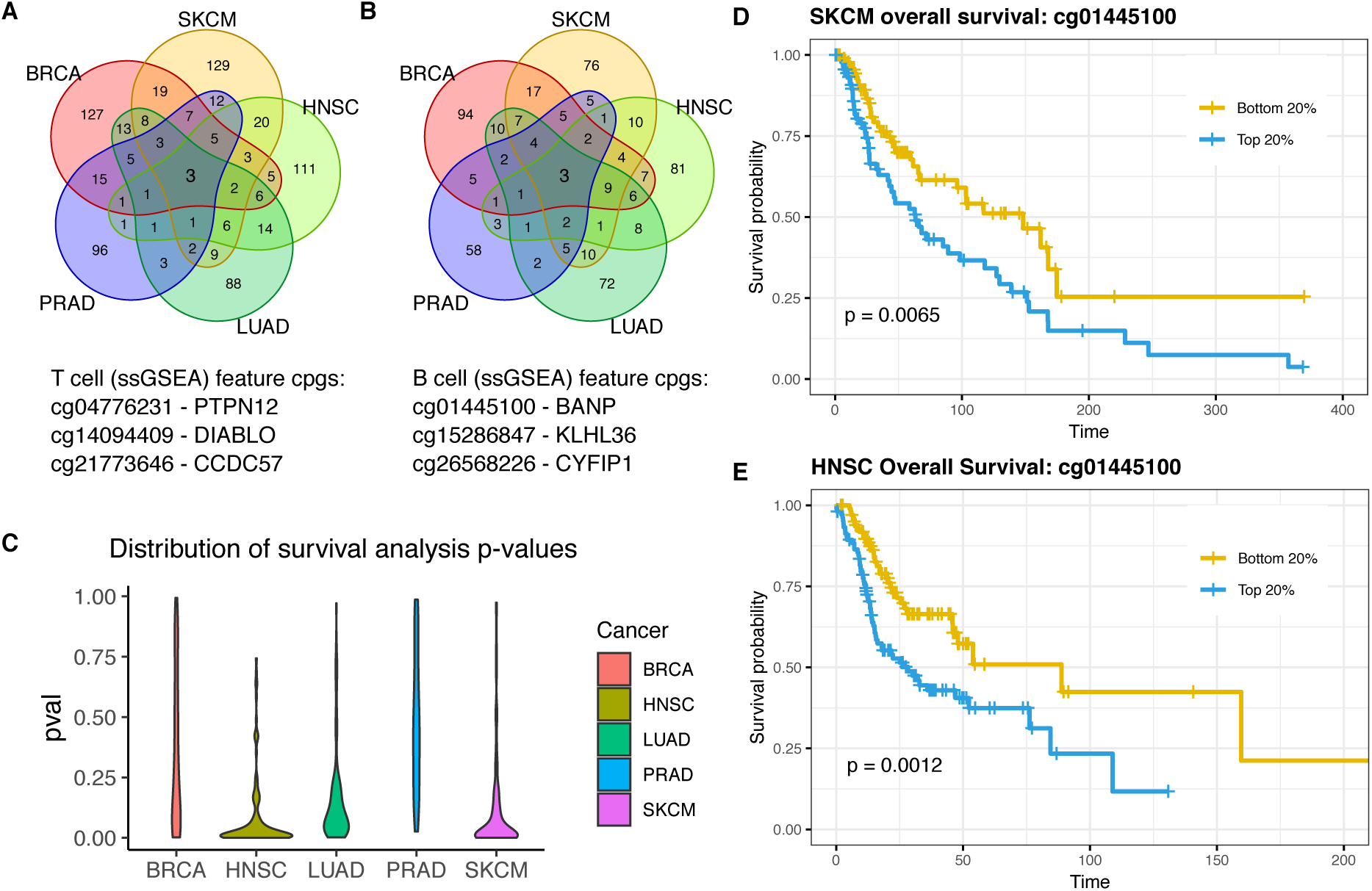
Methylation panels for estimating tumor infiltrated immune cell scores. **(A)** and **(B)** CpG sites shared by methylation panels of T cells and B cells in five cancer types, BRCA, SKCM, HNSC, LUAD, and PRAD. **(C)** Association between methylation and cancer prognosis varies by cancer type. SKCM and HNSC achieved the best association, whereas PRAD was the worst. **(D)** and **(E)** Examples of Kaplan-Meier curves of marker CpGs in our methylation panel. cg01445100 is located near to the promoter region of gene BANP. Lower methylation level of cg01445100 is associated with better prognosis in SKCM and HNSC.

In addition, we studied predictive CpG sites that are shared across different TIL cell types. We found the importance score of cg07638500 is greater than 0.9 in estimating B cells, T cells, cytotoxic cells as well as overall infiltration scores (TIS and IIS) in SKCM (Supplementary Figure 3). cg07638500 is located in the gene body of myosin light chain kinase (MYLK). Previous study has found that the gene expression level of MYLK is associated with the invasiveness of uveal melanoma cells [34]. The Venn diagrams of overlapping CpGs across cell types for all five cancer types are available in Supplementary Figure 3. These results on related functions of top methylation markers suggest that methylation array contains important molecular information of the tumor microenvironment that needs to be further investigated. We documented the methylation biomarkers with importance scores greater than 0.9 (methylation panel) for 33 immune infiltration scores across 10 cancer types in Supplementary Table 2. This will serve as an important resource of methylation biomarkers in understanding the tumor microenvironment.

As an important validation, we investigated the prognosis power of CpGs in our methylation panel using Kaplan-Meier curves and log-rank test. Tumor samples in the top 20th percentile of selected probe value were compared with those in the bottom 20th percentile. We found that the association between methylation and patient survival varies by cancer type (Figure 3C). Among ten cancer types in our analysis, SKCM showed the strongest association between CpG biomarkers and prognosis. For example, we found that lower methylation values of cg01445100 (BANP) is associated with improved survival (adjusted p-value <0.01) in SKCM as well as HNSC patients (Figure 3D and 3E). These results indicate that the investigation of DNA methylation signatures may provide novel insights to the understanding of tumor progression and prognosis.

### Biological functions of top methylation probes

We next investigated the biological functions of genes next to methylation probes in our methylation panel. Gene ontology analysis of the genes closest to selected methylation probes reveals enriched processes related to lymphocyte activation, signal transduction, and regulation of cell adhesion in many cancer types (see Supplementary Table 3 for GO results for BRCA, HNSC, HNSC, LUAD, PRAD and SKCM). The top five enriched processes in SKCM are small GTPase mediated signal transduction, lymphocyte activation, cellular component organization and cell surface receptor signaling pathway. The fraction of methylation marker probes in close proximity to immune marker genes [9] varies from 0 to 26% in all cancer types (Supplementary Table 4). We compared the predictive power of immune marker gene related CpGs to the predictive power of our methylation panel, and found a lower prediction accuracy using immune marker gene related CpGs (Supplementary Figure 4). This result suggests that some CpGs that are not in close proximity of immune marker genes also plays an important role in determining the tumor microenvironment.

**Figure 4.**
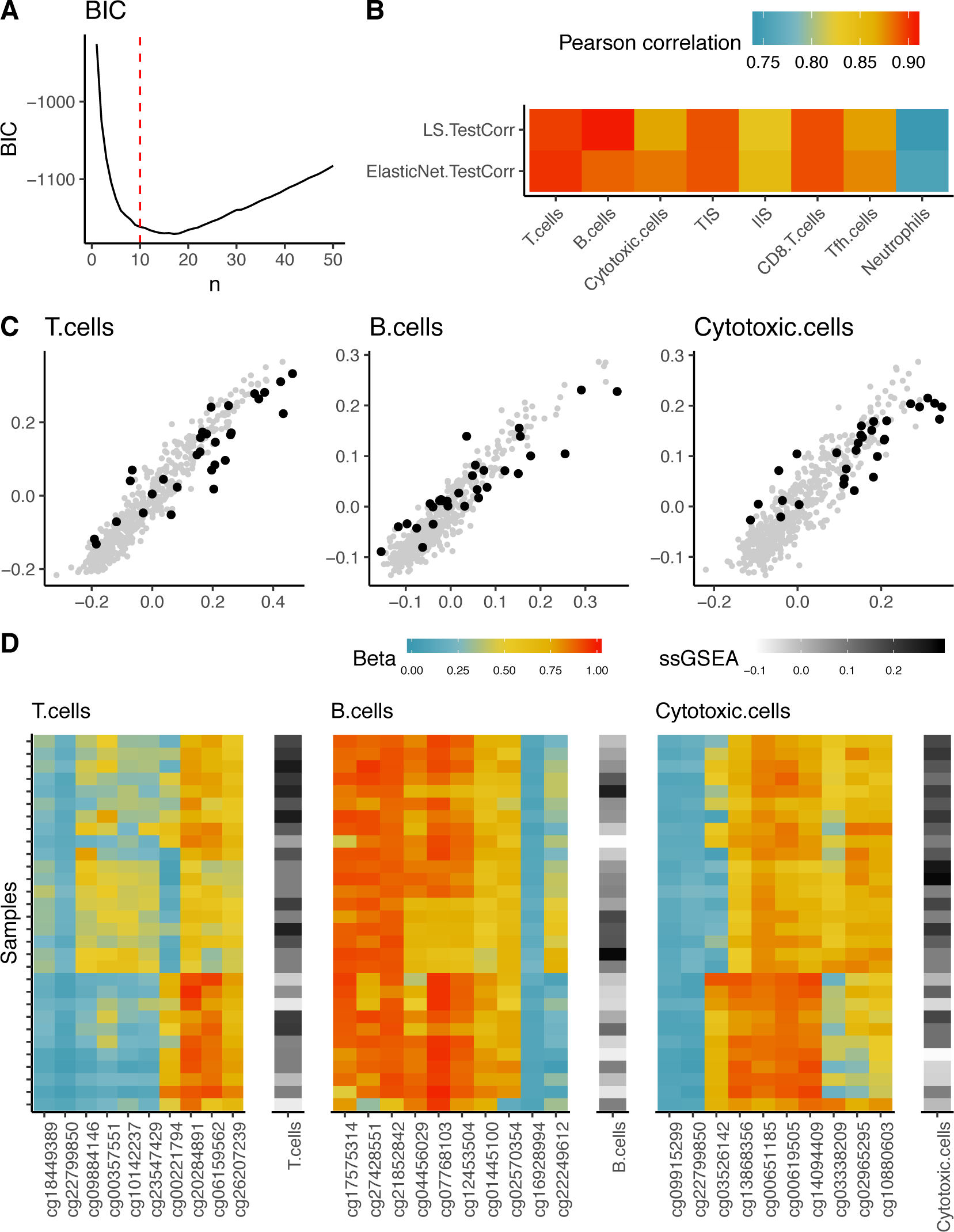
Validation of our regression model using 30 new melanoma samples. **(A)** BIC as a function of the number of features in forward stepwise selection of linear regression models. Red line indicates the final selected number of features. **(B)** Pearson correlation of gene expression predicted TIL scores with methylation predicted TIL scores using LS10 and ElasticNet model for 30 new melanoma samples. **(C)** Scatter plot of gene expression predicted TIL scores vs. methylation predicted TIL scores. Grey dots: TCGA skin cancer melanoma samples. Black dots: 30 new melanoma samples. **(D)** Heatmaps for methylation levels of features in RL10 model for 30 new melanoma samples. Cell types from left to the right: T cells, B cells, cytotoxic cells. ssGSEA scores are visualized on the right of each panel.

### Regression models for TIL estimation in SKCM

Our analysis of TCGA data shows that methylation profiles achieves the best prediction accuracy in SKCM. Next, we took two approaches to build practical tools for estimating immune cell infiltration scores in SKCM using our methylation panel. In this analysis, all TCGA samples were used as the training set. The first approach is to directly apply elastic net regression models (ElasticNet model) on the training set. Regularization parameters in the ElasticNet model were selected by cross-validation. The second approach is to use forward stepwise selection with linear regression models. We plotted model BIC as a function of the number of features in this model, and found that model BIC first dropped rapidly then gradually increased as more CpGs were included (Figure 4A). We chose the linear regression model with ten CpG sites as our final model (LR10 model), in order to keep the regression model easy to implement while retain high prediction accuracy. To further validate our predictive models, we performed EPIC 850K methylation array on 30 independent SKCM samples profiled with matched RNA-seq. A quantile normalization (see methods) was applied to calculate the beta values for EPIC samples. We found high concordance between ssGSEA and estimated TIL scores in EPIC sample using both ElasticNet model and LR10 model (Figure 4B and 4C). Importantly, no significant difference between prediction power in EPIC samples was observed between the ElasticNet and LR10 model. The heatmap of beta values of CpG sites in the LR10 model in our new melanoma samples reveals stratification of samples with high vs. low levels of immune cell infiltration (Figure 4D). These results further demonstrated the power of our predictive model in evaluating SKCM TIL abundance and the robustness of our method to experimental variations.

Here we present the LR10 model for estimating B cell, T cell and cytotoxic cell infiltration in SKCM using arcsin transformed methylation beta values, arcsin(2*beta-1). Coefficients were rounded to the third decimal places.

y[SKCM, B.cells] = 0.163 - 0.097 ×cg01445100 - 0.042 ×cg07768103 - 0.066 ×cg12453504 + 0.105 ×cg16928994 + 0.040 ×cg17575314 + 0.036 ×cg22249612 + 0.019 ×cg27428551 - 0.024 ×cg04456029 + 0.072 ×cg21852842 + 0.021 ×cg02570354

y[SKCM, T.cells] = 0.356 - 0.016 ×cg00221794 + 0.033 ×cg00357551 - 0.035 ×cg06159562 + 0.082 ×cg10142237 + 0.070 ×cg18449389 - 0.063 ×cg20284891 + 0.099 ×cg22799850 + 0.043 ×cg23547429 - 0.069 ×cg26207239 + 0.019 ×cg09884146

y[SKCM, Cytotoxic.cells] = 0.364 - 0.056 ×cg00619505 + 0.104 ×cg09915299 - 0.044 ×cg13868356 - 0.044 ×cg14094409 + 0.079 ×cg22799850 + 0.021 ×cg03338209 - 0.047 ×cg00651185 + 0.020 ×cg02965295 - 0.014 ×cg03526142 + 0.019 ×cg10880603

## DISCUSSION

Combining molecular data from both DNA and RNA will largely contribute to our understanding of the tumor-immune ecosystem and its association with clinical outcomes such as patient responses from receiving immune checkpoint inhibitors. Existing methods for tumor immune cell decomposition are largely based on gene expression data, while DNA data are only considered informative for tumor mutational and clonality analysis. Due to the high sensitivity of DNA methylation profiles to cell mixtures, digital dissection of the tumor immune landscape based on methylation data might be a promising new trend in cancer research. It has been previously demonstrated that tumor purity estimates from DNA methylation profiles (LUMP) showed high concordance with the estimates produced by other molecular data [35]. Compared to gene expression data, DNA methylation has further appealing features: (1) DNA methylation signature is generally more reliable and presumably more reproducible in routine analysis than gene expression. The addition of methyl group to cytosine is a very stable chemical alternation. RNA expression value, on the other hand, is complicated by many factors and is more likely to be affected by somatic copy number variations. Additionally, DNA itself is also chemically more stable than RNA. (2) Epigenetic elements have a high translational potential as these elements potentially represent the most “druggable” targets in cancer. For example, among all genes that are modulated in cancer cell treated with the FDA-approved drug 5-azacityidine, about twenty percent are related to immune regulation [ref]. The analysis procedure developed in this study will not only facilitate the secondary data analysis of retrospective data where both gene expression and DNA methylation data are available, but will also greatly aid future prospective studies in which only DNA methylation data are available. (3) Methylation arrays are less expensive compared to sequencing and flow cytometry methods. DNA methylation array-based assays are also forward compatible: most CpG sites in old platforms will be included in newer assays. This feature makes the further validation of our results possible with arrays such as the EPIC array or sequencing based methylation assays.

As previously discussed, methods for deconvolving cell content from molecular data fall into two main categories: reference-based and reference-free [36]. The reference-based method is motivated by the gene expression based deconvolution using constrains (e.g., CIBERSORT), which is implemented based on the cell-type-specific gene expression profiles (GEP). The default reference GEP used in CIBEERSORT is called LM22, which contains 547 genes for distinguishing 22 cell types. These genes were selected based on the differential expression analysis from expression profiles of purified cell subsets, and ideally, they should be exclusively expressed in each cell type they are representing. Similarly, a cell-type-specific methylation reference profiles can be constructed by analyzing methylation profiles of purified cell groups, using differential methylation (DM) analysis or analysis for identifying differential methylated regions (DMRs). On the other hand, reference-free methods do not require reference gene expression or methylation profiles, but use unsupervised methods such as surrogate variable analysis (SVA) to discover potential cell type groups. However, because reference-free methods do not provide direct estimate of the cell type fractions, they are only suitable in adjusting for confounding factors in EWAS and not applicable if the main purpose is to estimate the abundance of immune cells in solid tumor tissues. Both the recently developed MethylCIBERSORT and our procedure fall into the reference-based method category, despite the differences in constructing reference panel and the predictive model.

The biomarker discovery scheme developed in this study provides a practical solution for the problem of limited methylation data generated from FACS-purifed cells. An ideal reference panel should include CpGs that are exclusively highly-methylated or unmethylated in the cell type that they are representing. This is usually done by comparing DMRs from pairwise DM analysis from any two purified cell populations. Given the considerable heterogeneity in the tumor microenvironments and in the molecular profiles of malignant cells, we argue that the differential-methylation-based strategy should be applied separately in each cancer type. However, most of the existing molecular deconvoultion methods, including CIBERSORT and MethylCIBERSORT, have been built based on a unified panel from the same source of the purified immune cells. While there is a paucity of purified methylation libraries for each cancer type, the recent development in single cell DNA methylation analysis offers another possibility. Single cell methylation data allows us to isolate the cell populations in silico and built single cell reference methylation profiles (sc-MPs), similar to the concept of the single-cell reference GEP (sc-GEP). Our model-based scheme can be adapted to build sc-MPs in the future. Another promising extension of our method is to select CpGs that are associated with immunotherapy responses. Immunotherapy, such as immune checkpoint blockade (ICB), has rapidly become a first-line treatment option in many cancer types. There are yet no methylation biomarkers available to predict response to ICBs.

